# Directed migration shapes cooperation in spatial ecological public goods games

**DOI:** 10.1101/577205

**Authors:** Felix Funk, Christoph Hauert

## Abstract

From the microscopic to the macroscopic level, biological life exhibits directed migration in response to environmental conditions. Chemotaxis enables microbes to sense and move towards nutrient-rich regions or to avoid toxic ones. Socio-economic factors drive human populations from rural to urban areas. However, migration affects the quantity and quality of desirable resources. The effect of collective movement is especially significant when in response to the generation of public goods. Microbial communities can, for instance, alter their environment through the secretion of extracellular substances. Some substances provide antibiotic-resistance, others provide access to nutrients or promote motility. However, in all cases the maintenance of such public goods requires costly cooperation and is consequently susceptible to exploitation. The threat of exploitation becomes even more acute with motile individuals as defectors can avoid the consequences of their cheating.

Here, we propose a model to investigate the effects of targeted migration based on the production of ecological public goods and analyze the interplay between social conflicts and migration. In particular, individuals can locate attractive regions by moving towards higher cooperator densities or avoid unattractive regions by moving away from defectors. Both migration patterns not only shape an individual’s immediate environment but also affects the population as a whole. For example, defectors hunting cooperators in search of the public good have a homogenizing effect on population densities. They limit the production of the public good and hence inhibit the growth of the population. In contrast, aggregating cooperators promote the spontaneous formation of heterogeneous density distributions. The positive feedback between cooperator aggregation and public goods production, however, poses analytical and numerical challenges due to its tendency to develop discontinuous distributions. Thus, different modes of directed migration bear the potential to enhance or inhibit the emergence of complex and sometimes dynamic spatial arrangements. Interestingly, whenever patterns emerge in the form of heterogeneous density distributions, cooperation is promoted, on average, population densities rise, and the risk of extinction is reduced.

**Author summary:** The production and maintenance of shared environmental resources such as access to nutrients in microbial communities or potable water in human societies require the cooperation of groups of individuals. However, cooperation is costly and prone to exploitation. If too many individuals follow selfish interests and spoil their environment, the group and possibly the entire population suffers. Nevertheless, many forms of biological life – from humans to microbes – migrate in response to resource availability. Here, we analyze the interplay of the social conflict in public goods production and targeted migration. In particular, we find that aggregation of cooperators can enhance or trigger the spontaneous formation of heterogeneous spatial distributions, which promote cooperation and result in higher population densities. Conversely, attempts to avoid defectors increases the risk of extinction because it tends to homogenize population distributions and lower population densities.

## Introduction

Directed migration is a phenomenon commonly observed in nature. Microbial populations such as *Escherichia coli* actively seek areas with higher concentrations of substances, such as amino-acids, through chemotaxis [1] and avoid toxic regions [2]. Birds sense seasonal changes and migrate to more hospitable climes [3]. Socio-economic factors attract people from rural communities to cities [4]. In all cases, individuals migrate to improve their access to resources.

Environmental factors influence the abundance of resources, but some resources such as public goods are produced and maintained by the population itself and require cooperation. Prominent examples are drinking water, clean air or the climate on a global scale [5] as well as the production of extracellular substances to access food [6] or to grant resistance to antibiotics [7, 8] on a microscopic scale. However, cooperation is costly and threatened by non-cooperating defectors [9]. Defectors are free-riders exploiting public goods without contributing themselves and hence the public resources dwindle whenever defection becomes common.

Ecological public goods introduce ecological dynamics into evolutionary games through variable population densities based on the production of the public good [10]. Thus, defectors cause decreases in population densities, which results in smaller interaction group sizes. In sufficiently small groups cooperation may become dominant and the population can recover. This feedback between population densities and interaction group sizes enables cooperators and defectors to coexist. In particular, this includes the most interesting case where public goods production is crucial for the survival of the population [11]. This represent our baseline scario here.

In a spatial context, undirected movement introduces rich dynamics including the spontaneous emergence of heterogeneous density distributions resulting in pattern formation and chaotic spatio-temporal dynamics [12]. The pattern formation is driven by Turing instabilities [13], which have also been reported for purely ecological models such as predator-prey or chemotaxis models [14, 15]. Whenever patterns form, populations thrive through an increase in cooperation, raised population density, and reduced risk of extinction. However, spatio-temporal chaos and pattern formation also causes spatial heterogeneity in the abundance and availability of the public good, which naturally leaves some locations more attractive than others.

Through directed migration, cooperators and defectors can improve access to the public good by actively seeking cooperation or reduce exploitation by avoiding defection. Here, we extend the spatial ecological public goods game [12] by incorporating targeted migration in the “selection-diffusion” system, and analyze how directed migration shapes cooperation, and pattern formation, in particular. Based on the stability analysis of the dynamical equations and supported by numerical integration, we derive detailed criteria for pattern formation for the different modes of migration.

Directed migration can trigger, enhance or inhibit the spontaneous emergence of steady, quasi-steady or dynamic heterogeneous density distributions. Somewhat surprisingly, cooperators fleeing from defectors homogenize the density distribution, and endanger the population whenever cooperators spread themselves too thin. In contrast, if defectors avoid each other, cooperators can accumulate and patterns emerge. This allows cooperation to thrive under conditions, in which neither a well-mixed population nor a diffusing migration would survive. As expected, aggregating cooperators also facilitate pattern formation through positive feedback between cooperator densities and cooperator migration: cooperator aggregation gives rise to steeper gradients of the public good and thus creates even stronger incentives to migrate in towards those locations. However, the positive feedback comes with analytical and numerical challenges due to the potential to give rise to discontinuous distributions. Through the promotion or inhibition of spatial heterogeneity, directed migration is a critical determinant for cooperation, the survival of the population, and conducive to promoting and maintaining diversity of ecosystems.

### Ecological public goods

In traditional public goods games *N* individuals gather in a group and each individual chooses to cooperate and invest into a common pool at a cost *c*, or to defect and shirk the investment. The common pool is multiplied by a factor *r >* 1 and equally divided among all individuals within the group. This results in a payoff of *P*_*D*_ = *n_C_ ⋅ rc/N* for defectors and *P_C_* = *P*_*D*_ − (1 *− r/N*)*c* for cooperators when facing *n_C_* cooperators among the *N −* 1 other group members. For *r < N* every participant is tempted to withhold their investment because *P*_*C*_ < *P*_*D*_ and hence to forego the benefits of the public good to the detriment of all. Had everyone cooperated the payoff would be (*r* − 1)*c* > 0. This conflict of interest between the group and the individual constitutes the social dilemma at the heart of public goods interactions [16–18]. Only for *r > N* these interests align.

In well-mixed populations with normalized densities (frequencies) of cooperators, *u*, and defectors, *v* = 1 *− u*, both strategies encounter, on average, *n*_*C*_ = *u*(*N −* 1) cooperators among their interaction partners in randomly assembled interaction groups. In evolutionary game theory payoffs represent fitness and hence determine the rate of change of the population composition as captured by the replicator dynamics *∂_t_u* = *u*(1 *− u*)(*P_C_ − P_D_*), where *∂_t_* denotes the time derivative [19]. Consequently, cooperation dwindles, *∂_t_u <* 0, whenever cooperators obtain lower payoffs in the public goods game than defectors, i.e. if *P_C_ − P_D_* = *−c*(1 *− r/N*) *<* 0. Indeed, for *r < N* defection dominates and *∂_t_u <* 0 always holds. Conversely, for *r > N* cooperation spreads in the population, *∂_t_u >* 0.

The replicator equation does not take ecological quantities such as variable population densities into account. In ecological systems, the presence of the public good promotes population growth through improved access to food resources [6] or increased resistance to environmental threats such as antibiotics [7]. The interplay between ecological and evolutionary dynamics in such ecological public goods interactions is captured by

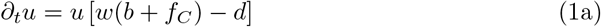

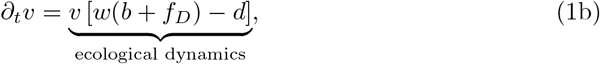

where *w* = 1 − *u* − *v* reflects reproductive opportunities that diminish for increasing population densities [11]. Selection is based on ecological public goods interactions affecting the birth rates of cooperators and defectors through their average payoffs *f*_*C*_ and *f*_*D*_, respectively, while the death rate is constant, *d*. More specifically, the effective birthrates, *w*(*b* + *f*_*i*_), are determined by

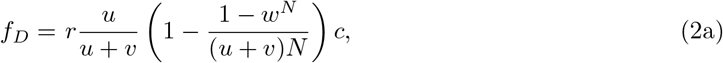

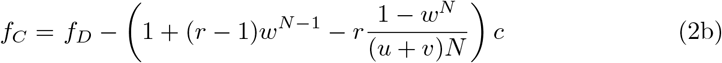

plus the baseline birthrate, *b*, because in the limit *v →* 1 the payoff *f*_*C*_ can become negative, which is biologically not meaningful [11]. The ecological dynamics results in variable interaction group sizes with an effective, expected group size of *S* = (*u* + *v*)*N*. Thus, as long as *S > r*, defection dominates, the public resource is overexploited and the population density declines. Consequently, interactions occur in smaller groups and returns are split among fewer individuals. For sufficiently small groups, *S < r*, cooperation becomes favourable again, the population density recovers, *S* increases and the cycle continues.

Even in the absence of spatial dimensions, rich dynamics are observed especially under environmental stress, *d > b*, where the survival of the population hinges on the availability of the public good [10]. Varying population densities enable the coexistence of cooperators and defectors. More specifically, an interior equilibrium *Q* undergoes a Hopf-bifurcation [20] when increasing the rate of return of the public good, *r*, giving rise to stable and unstable limit cycles [11]. For *r < r*_Hopf_, *Q* is unstable, and well-mixed populations are doomed. However, for *r > r*_Hopf_, the equilibrium *Q* allows cooperators and defectors to coexist but the basin of attraction of the equilibrium *Q* remains limited. More specifically, if cooperators are too rare for the public goods production to offset the death rate *d*, or if defectors abound, the population may still go extinct.

Here we focus on the case were the death rate exceeds the baseline birthrate, *d > b*, such that only the production of the public good can prevent extinction. This scenario can be interpreted as a microbial population in a biocide where the multiplication factor *r* reflects the effectivity of the public good to oppose the detrimental effect of the toxic environment. The following extensions explore the effects of different modes of motility on the public goods production and survival of the population.

## Materials and methods

### Spatial ecological public goods

The spatial dynamics of ecological public good interactions with undirected (diffusive) migration can be formulated as a selection-diffusion process

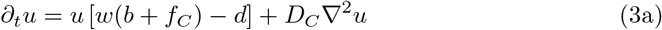

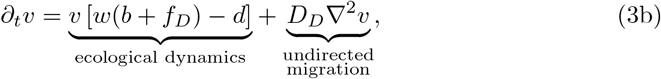

where the diffusion constants *D*_*C*_ and *D*_*D*_ reflect the migration of cooperators and defectors, respectively [12]. In spatial settings, undirected migration promotes coexistence of cooperators and defectors even if the coexistence equilibrium *Q* is unstable. In particular, if defectors run faster than cooperators, *D*_*D*_ > *D*_*C*_, the dynamics exhibit Turing instabilities, where cooperators serve as activators and defectors as inhibitors. This enables the population to survive through spontaneous pattern formation. Close to the Hopf-bifurcation with *r < r*_Hopf_ (*Q* unstable), the influence of temporal oscillations gives rise to chaotic spatio-temporal dynamics [21], which facilitate population survival even for equal diffusion rates, *D*_*D*_ = *D*_*C*_. Conversely, for slower diffusion of defectors, *D_D_ < D_C_*, activation through cooperation is no longer tenable, spatial effects disappear and either results in homogeneous coexistence for *r > r*_Hopf_ (*Q* stable) or extinction for *r < r*_Hopf_. This implies that attempts of cooperators at outrunning defectors are futile.

Here, we extend the spatial ecological public goods model, Eq. (3), by introducing *directed* migration, which enables cooperators and defectors to bias their movement towards more attractive regions.

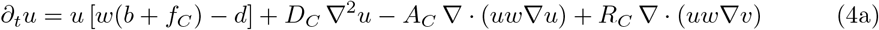

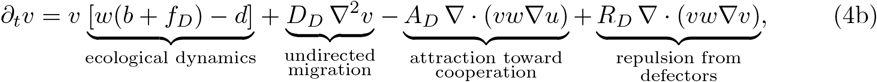

with *A*_*C*_, *A*_*D*_, *R*_*C*_, *R*_*D*_ ≥ 0 and where terms of the form *−K∇ ⋅* (*φw∇ψ*) reflect that individuals of type *φ* are attracted to the gradient of type *ψ* proportional to reproductive opportunities, *w*, at a non-negative rate *K* (for a detailed microscopic derivation see S1 Appendix). The density of cooperators directly translates into the rate of production of public goods and hence serves as a proxy for its availability and the quality of the environment. For example, the term *−A_D_∇ ⋅* (*vw∇u*) reflects *hunting defectors* in search of public goods that are attracted to higher densities of cooperators at a rate *A*_*D*_. Note that the negative sign indicates movement toward higher densities. Similarly, *−A_C_ ∇ ⋅* (*uw∇u*) represents *aggregating cooperators* that are attracted to their kind. In contrast, the density of defectors serves as a proxy to avoid exploitation and reduce competition. Instead of directly sensing the density of defectors their excretions from public goods consumption could reflect the poor quality of the environment. Thus, *R*_*C*_ ∇ ⋅ (*uw∇v*) reflects *fleeing cooperators* that avoid higher densities of defectors at a rate *R*_*C*_, whereas *R*_*D*_∇ ⋅ (*vw∇v*) refers to *spreading defectors* that steer clear of their kind.

Rich dynamics unfold under directed migration as showcased in Fig. 1, ranging from the spontaneous formation of quasi-stable or stable patterns (Fig. 1A) and ever-changing, chaotic spatio-temporal dynamics (Fig. 1B) to cooperators aggregating under a self-reinforcing migration response (positive feedback, Fig. 1C).

**Fig 1.**
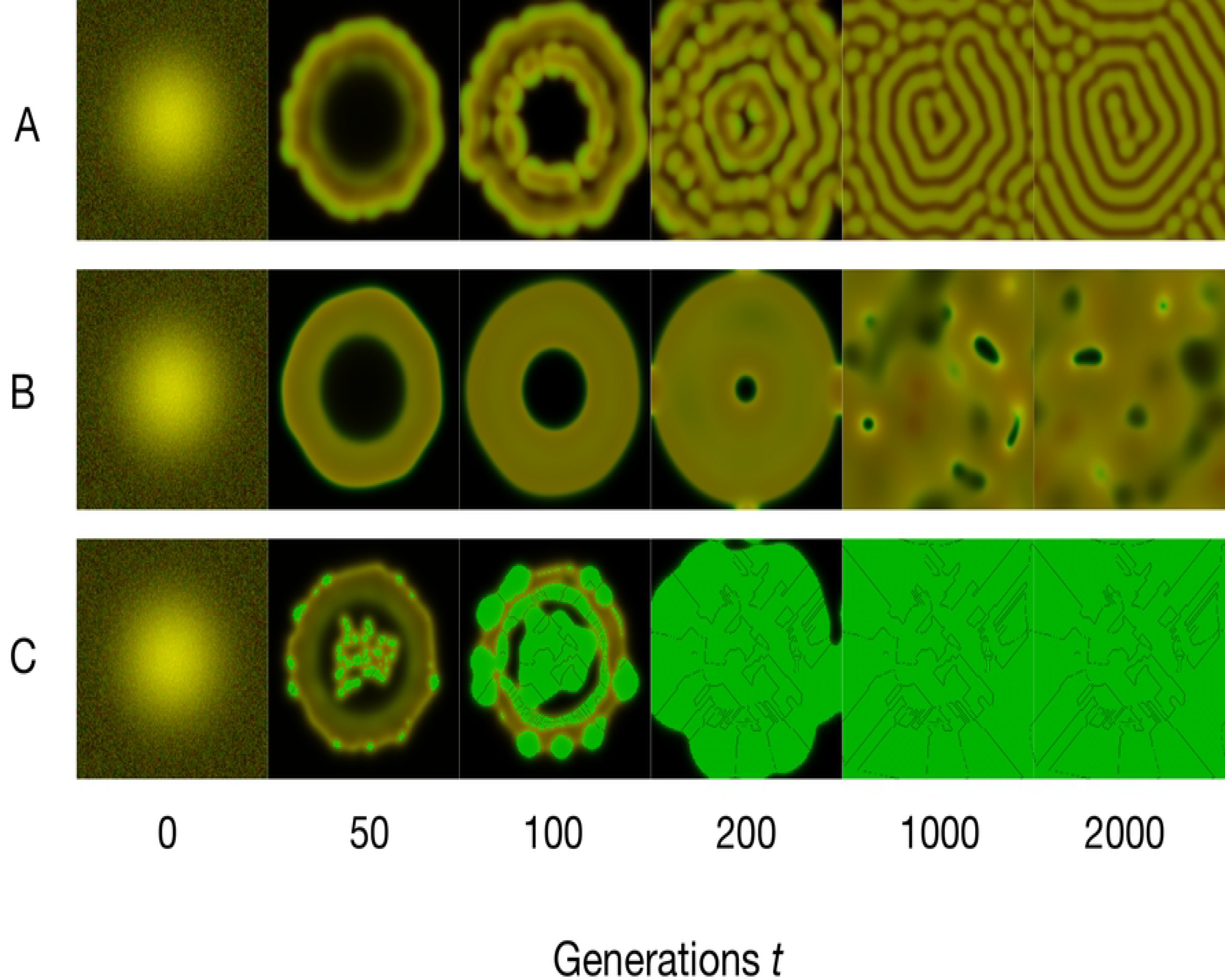
Pattern formation under directed migration. Snapshots of the density distribution of cooperators (green) and defectors (red) across the periodic *L × L* domain after *t* = 0, 50, 100, 200, 1000, 2000 generations. The color brightness indicates the density with coexistence (yellow) and vacant space (black). (A) Spreading defectors spontaneously form and settle in quasi-steady patterns (*R*_*D*_ = 8). (B) Chaotic spatio-temporal dynamics unfold for hunting defectors (*A*_*D*_ = 4). (C) Aggregating cooperators induce positive feedback: as more cooperator leave scarcely populated areas in favor of highly cooperative locations, the divide between neighbouring areas grows and discontinuities can develop at the interfaces. (*A*_*C*_ = 1) *Parameters:* public goods interaction: *b* = 1*, r* = 2.325*, d* = 1.2*, N* = 8*, c* = 1; diffusion: *D*_*C*_ = *D*_*D*_ = 0.1; directed movement: *A*_*C*_ = *A*_*D*_ = *R*_*C*_ = *R*_*D*_ = 0, unless otherwise indicated; discretization in space and time: *L* = 75*, dx* = 0.375*, dt* = 0.01; noisy Gaussian initial condition: *u*(*x, y*) = *v*(*x, y*) = 1*/*5 *⋅* exp[*−*((*x − L/*2)^2^ + (*y − L/*2)^2^)*/*16^2^].

## Results

An overview of the long-term effects of the four types of directed migration on spatial ecological public goods interactions is summarized in Fig. 2. Each type has distinct effects on pattern formation and population survival. For *r < r*_Hopf_, aggregating cooperators, *A*_*C*_, (see Fig. 2A) and spreading defectors, *R*_*D*_, (see Fig. 2D) increase the chances of survival by promoting spontaneous formation of heterogeneous density distributions. For aggregating cooperators, this process is facilitated by a slower rate of expansion. In particular, aggregating cooperators directly oppose the effects of cooperator diffusion *D*_*C*_ and can even bring population expansion to a halt (see panel A in S2 Figure). In fact, sufficiently high *A*_*C*_ triggers a positive feedback between cooperator densities (or public goods production) and aggregation, which pose analytical and numerical challenges discussed below. In contrast, hunting defectors, *A*_*D*_, increase competition in areas where cooperators aggregate (see Fig. 2C) and fleeing cooperators, *R*_*C*_, explore space to avoid exploitation (see Fig. 2B). In both cases cooperator densities and hence the concentration of the public good are evened out. This suppresses pattern formation and renders the population more prone to extinction through global temporal fluctuations. For *r > r*_Hopf_, coexistence of cooperators and defectors in well-mixed populations is stable and the spatial patterns tend to fade away. The only exception is the aggregation of cooperators, which may prevent expansion and hence colonization of empty territory.

**Fig 2.**
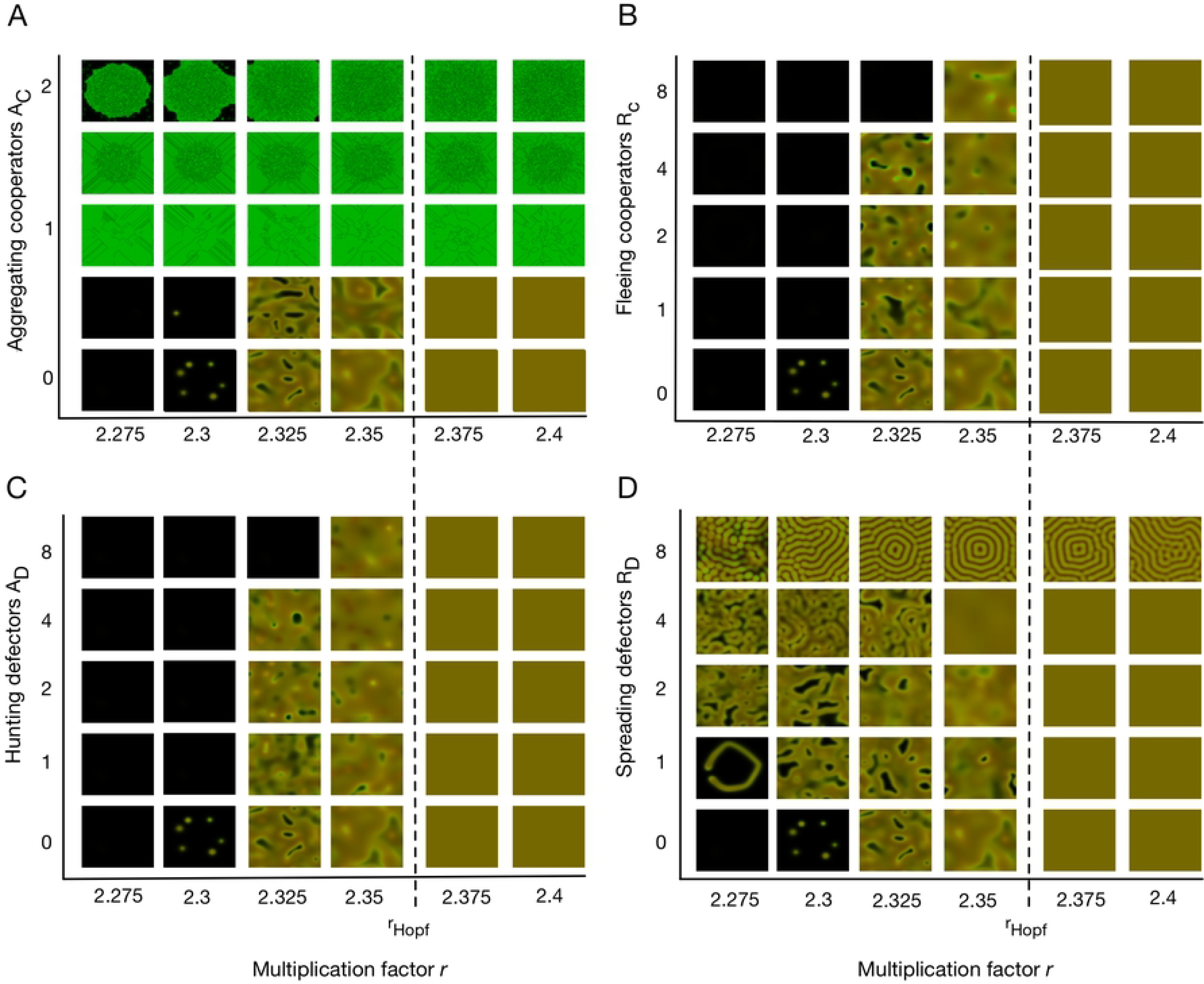
Spatial dynamics for the four types of directed migration. Each of the four panels (A)-(D) shows snapshots of the spatial distributions after *t* = 2000 generations as a function of the multiplication factor *r* when varying one of the four modes of migration. The bottom row shows the dynamics of purely diffusive migration in the proximity of *r*_Hopf_ ≈ 2.365 as a reference. (A) Aggregating cooperators oppose the effects of diffusion and hence slow down the population expansion. The positive feedback between cooperator densities and aggregation can cause discontinuous distributions and the breakdown of numerical methods. (B) Hunting defectors increase competition in areas where cooperators concentrate. This allows cooperators to escape into unpopulated terrain but also increases the risk of extinction. (C) Fleeing cooperators avoid defectors and hence readily explore vacant space but with the defectors at their tails also tend to spread themselves too thin and risk extinction. (D) Spreading defectors reduce exploitation on cooperator aggregates and consequently promote pattern formation as well as population survival. *Parameters* as in Fig. 1 unless otherwise indicated.

### Emerging patterns

Public goods production and population survival are both crucially linked to the spontaneous emergence of spatial patterns. Here we derive necessary conditions for the onset of pattern formation. An analytical understanding of the pattern formation process is obtained by considering homogeneous population densities reflecting the coexistence equilibrium *Q* of the well-mixed population dynamics, Eq. (1). A small perturbation *p*(*t, x, y*) = *ϵ* exp(*ik*(*x* + *y*) *− t*) of mode *k*, where *k* = 0 reflects the temporal mode and *k >* 0 spatial modes, may get amplified by the dynamics and give rise to the emergence of heterogeneous density distributions, also called Turing patterns [13]. The linearized dynamics in the vicinity of *Q* = (*u*_eq_*, v*_eq_*, w*_eq_) is given by

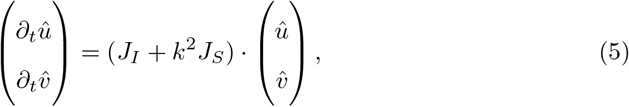

where 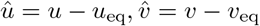 and *J*_*I*_, *J*_*S*_ represent the Jacobians from public goods interactions and spatial migration (directed and undirected), respectively:

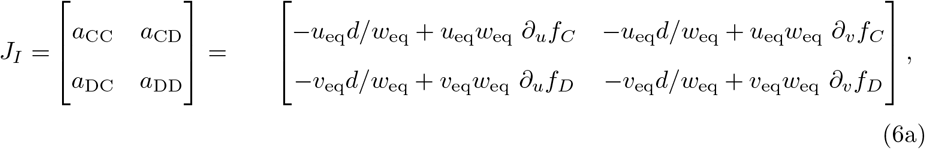

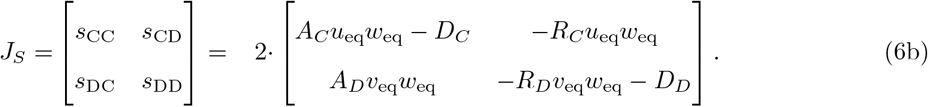

The largest eigenvalue (real part) of *J*_*I*_ + *k*^2^ *J*_*S*_ is a function of *k* and is called the dispersion relation *λ*(*k*) (see S2 Appendix). Modes with *λ*(*k*) *>* 0 are unstable indicating the potential for spatial patterns to emerge. If the spatial Jacobian *J*_*S*_ admits eigenvalues with positive real parts then *λ*(*k*) *>* 0 holds for all sufficiently large *k*, i.e. all spatial perturbations of sufficiently high frequency are unstable. This is not meaningful in natural systems. In order to ensure *λ*(*k*) *<* 0 for *k → ∞*, the condition det(*J*_*S*_) *>* 0 or, more precisely, *D_C_ > A_C_ u*_eq_*w*_eq_ is required, which means that at the homogeneous equilibrium, the effects of diffusion (undirected migration) need to outweigh effects of cooperator aggregation.

In spite of the above restrictions, intermediate modes can nevertheless become unstable through a combination of selection and migration. A necessary condition for pattern formation is

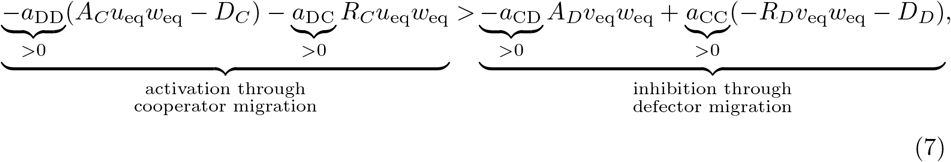

see S2 Appendix for details. Consequently, faster aggregating cooperators or spreading defectors (increased *A_C_, R_D_*) or slower undirected cooperator migration (decreased *D*_*C*_) promote the development of heterogeneous patterns by actively or passively promoting cooperator aggregation. In contrast, faster directed migration of defectors towards cooperation or of cooperators away from defection (increased *A_D_, R_C_*) as well as slower undirected defector migration (decreased *D*_*D*_) all increase exploitation of the public good, reduce the efficacy of cooperator aggregation and hence suppress pattern formation. Fig. 3 depicts the competing effects of activating and inhibiting forms of directed migration. The most unstable mode determines the characteristic length scale of the emerging spatial patterns (see S2 Appendix).

Hunting defectors, *A*_*D*_, and fleeing cooperators, *R*_*C*_, exhibit similar homogenizing effects. In contrast, significant qualitative differences arise in the promotion of pattern formation due to aggregating cooperators, *A*_*C*_, and spreading defectors, *R*_*D*_, in the sense that aggregating cooperators have the potential to fully suppress defection. In this case neither increases in *A*_*D*_ nor in *R*_*C*_ are capable of counteracting the positive feedback induced by *A*_*C*_ to recover coexistence.

**Fig 3.**
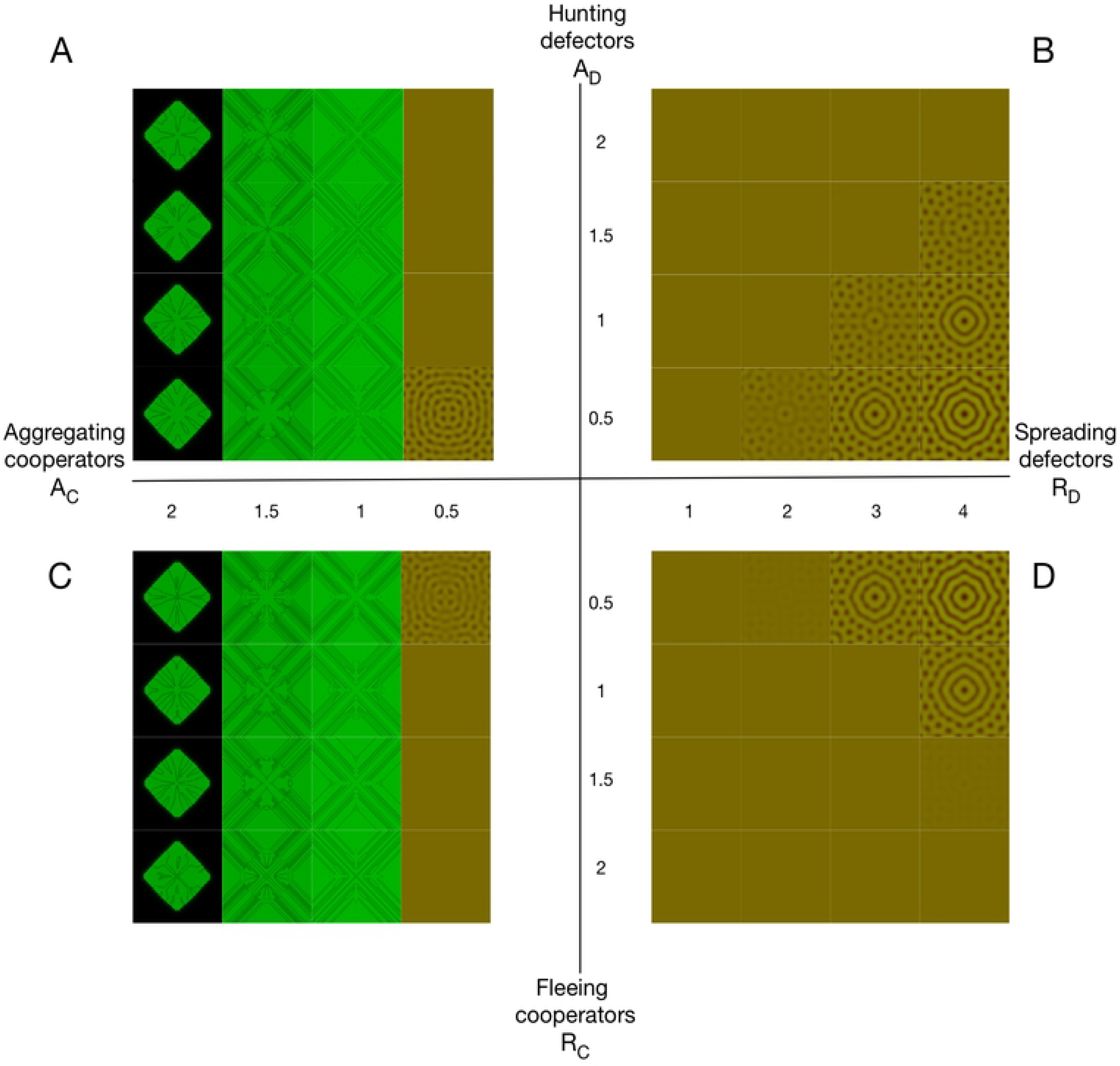
Pattern formation under competing forms of directed migration. (A), (C) Cooperator aggregation results in smooth patterns for small *A*_*C*_, which are suppressed by increasing *A*_*D*_, *R*_*C*_. However, for larger *A*_*C*_ the positive feedback between aggregation and densities of cooperators dominates. This eliminates defectors and gives rise to discontinuous distributions. Consequently, directed migration triggered by defector densities through *A*_*D*_ or *R*_*C*_ no longer matters. (B), (D) Spreading defectors, *R*_*D*_, also promote pattern formation but maintain smooth patterns, while increases in *A*_*D*_, *R*_*C*_ again suppress them. *Parameters:* Multiplication factor *r* = 2.4, *D*_*D*_ = 0.5 *> D_C_* = 0.1 to observe smooth patterns for *A*_*C*_; snapshots taken at *t* = 2000; Gaussian initial configuration without noise. Other parameters as in Fig. 1.

### Aggregating cooperators

The positive feedback between cooperator densities and cooperator migration is intrinsic to aggregating cooperators. Regions of higher cooperator densities attract cooperators, which further increases their densities and, more importantly, the gradient along the periphery. As a result the region exerts an even stronger attraction. In fact, for sufficiently large *A*_*C*_ the attraction prevents cooperators from exploring the available space thus the population remains localized. The potential of aggregating cooperators to give rise to discontinuous distributions through positive feedback lies at the core of the necessary condition that diffusion must outweigh aggregation at the homogeneous equilibrium: *D_C_ > A_C_ u*_eq_*w*_eq_. However, this condition is conservative and no longer sufficient for heterogeneous distributions. The gradient in cooperator densities, *∇u*, promotes further aggregation of cooperators despite being limited by reproductive opportunities, *w*, see Eq. (4). As a consequence the gradient further increases and discontinuities in the density distribution can develop. Interestingly, this effect turns out to be strong enough to completely suppress defection (c.f. Fig. 3). However, increasing gradients require finer discretization to numerically integrate the selection-migration dynamics, Eq. (4), and hence the emerging distributions depend on the discretization. The manifestation of discontinuous distributions becomes increasingly likely for larger values of *A*_*C*_ (see S3 Figure) but can also be triggered by pattern formation or large gradients in initial distributions. Increased and differing diffusion rates, *D_D_ > D_C_*, or spreading defectors, *R*_*D*_, lower the threshold for pattern formation and are additionally required to reliably observe smooth patterns through aggregating cooperators, *A*_*C*_.

### Social dynamics

Directed migration not only promotes or inhibits spontaneous pattern formation but also affects the frequency of cooperators as well as the population density and hence the survival of the population. Whenever patterns develop, cooperation is promoted and population densities consequently rise, see Fig. 4.

**Fig 4.**
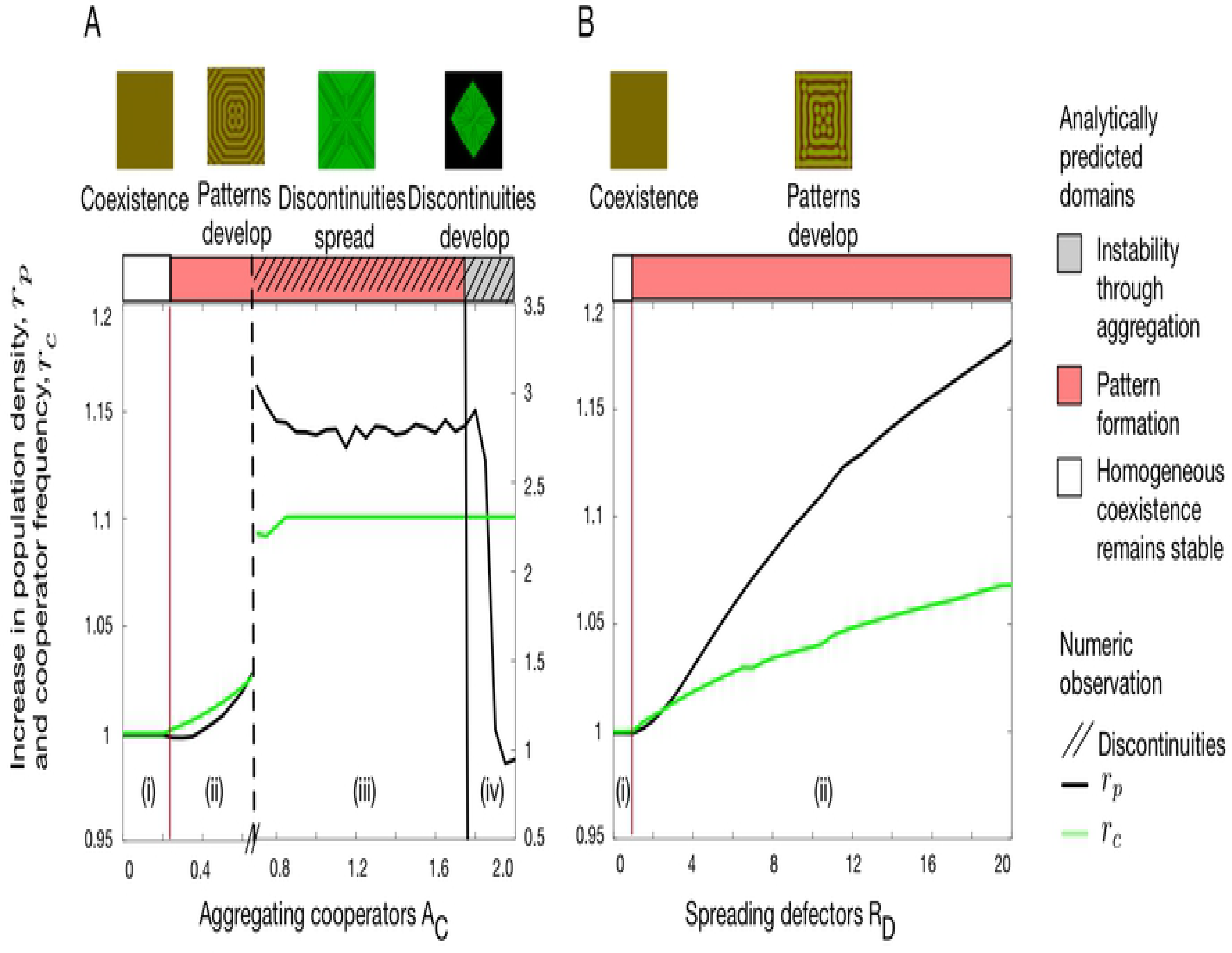
Effects of directed migration on population densities and frequency of cooperation as compared to unstructured populations. In the absence of directed migration (undirected migration only) homogeneous coexistence results with densities *Q* = (*u*_eq_*, v*_eq_). The two panels depict the ratio of the average spatial population density to the population density in unstructured populations, 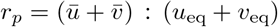 (black solid line), and the ratio of the average cooperator frequency to their frequency in unstructured populations, 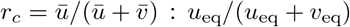 (green solid line) for (A) cooperator aggregation, *A*_*C*_, and (B) spreading defectors, *R*_*D*_. (A) increasing *A*_*C*_ gives rise to four dynamical regimes: (i) homogeneity maintained, (ii) formation of smooth patterns, (iii) discontinuities emerge, and (iv) population expansion prevented. (B) increasing *R*_*D*_ only results in two regimes: (i) homogeneity (ii) formation of smooth patterns. In order to facilitate comparisons of regions (i) and (ii) across panels the large effects of discontinuities in (iii) and (iv) refer to the scale on the right of panel (A). In either case, heterogeneous distributions increase (average) population densities and cooperator frequencies. In (A), once discontinuities develop, populations consist exclusively of cooperators. The decrease in *r*_*p*_ for large *A*_*C*_ relates to the fact that populations are unable to expand and hence the ratio depends on the initial configuration. In contrast, *r*_*c*_ remains unaffected because defectors are absent and cooperators are at the maximum frequency. *Parameters* as in Fig. 3.

This effect extends to multiplication factors below *r*_Hopf_, which cannot sustain unstructured populations. In this domain, undirected migration is capable of inducing spatial heterogeneity for *D_D_ > D_C_* and thereby promotes the survival of the population. Aggregating cooperators, *A*_*C*_, and spreading defectors, *R*_*D*_, further promote pattern formation and population survival even under conditions such as *D_C_ ≥ D_D_*, where undirected migration alone is unable to support the population, c.f. Eq. (7) and Fig. 2A, D. Moreover, the positive feedback of aggregating cooperators coupled with limited reproductive opportunities, offers the intriguing prospect that defectors can potentially get crowded out and driven to extinction, see Fig. 2A and Fig. 4A.

## Discussion

Cooperation is doomed in traditional public goods games – as paraphrased by Hardin’s “Tragedy of the Commons” [9]. In contrast, in ecological public goods interactions cooperators and defectors can coexist through the feedback between variable population densities and interaction group sizes [10] even if the reproductive performance and survival of the population hinges on the availability of public goods.

The introduction of continuous spatial dimensions allows population densities and social composition to vary across the domain. For undirected migration the ratio between the rates of cooperator and defector migration is crucial [12]. For *D_C_ > D_D_*, the population either settles in a homogeneous state with densities corresponding to the well-mixed coexistence equilibrium *Q*, if it is stable (*r > r*_Hopf_), or goes extinct if *Q* is unstable (*r < r*_Hopf_). In either case, attempts of cooperators at avoiding exploitation by outrunning defectors are futile. For *D*_*C*_ = *D*_*D*_, chaotic spatio-temporal dynamics enable the population to survive for *r* slightly below *r*_Hopf_ [21]. However, for *D_C_ < D_D_* cooperation is promoted through Turing instabilities that result in quasi-stable or dynamical patterns that increase cooperator frequency, overall population density, and enhance population survival.

Through undirected migration heterogenous density distributions of cooperators and defectors spontaneously arise. Consequently the abundance of public goods varies across space and renders some regions more attractive than others. This provides a natural incentive for directed migration – either to avoid poor areas or to seek out better ones. For example, chemotactic bacteria aggregate in patches in response to excreted attractants [22] or try to escape oxidative stress [2]. Instead of explicitly modelling the concentration of public goods and its waste products to assess the quality of the environment, we use the density of cooperators and defectors as proxies for the availability of public goods and the degree of exploitation, respectively. Thus, seeking cooperation and avoiding defection allows individuals to increase their access to public goods and improve their reproductive potential. Interestingly, even though the two migration patterns appear very similar, they may actually trigger movements in opposite directions. For example, cooperators that seek their kind aggregate at the centre of cooperative areas. The resulting increase in public goods also benefits defectors in that same location and may attract more defectors, which increases competition and exploitation. In contrast, cooperators that avoid defectors tend to migrate towards the periphery of cooperative areas and thereby effectively counteract cooperator aggregation. Analogous arguments apply to defectors seeking cooperators or avoiding their kind, respectively, but with opposite effects. More specifically, spreading defectors indirectly support cooperator aggregation by creating (temporary) refuges with low exploitation.

As a consequence, directed migration can both enhance as well as inhibit pattern formation. More specifically, aggregating cooperators or spreading defectors both promote or even trigger pattern formation. In particular, this extends into unfavourable parameter regions that otherwise result in extinction, such as when cooperators outpace defector, *D_C_ ≥ D_D_*, or small multiplication factors *r*, see Fig. 2A,D. Conversely, fleeing cooperators and hunting defectors suppress pattern formation by levelling out population densities. For *r < r*_Hopf_ this increases the risk of extinction. The complementary effects of the different modes of directed migration are captured in Eq. (7), which provides an analytical threshold for the onset of pattern formation. Regardless of whether spatial heterogeneities arise through directed or undirected migration, they invariably increase both the average frequency of cooperators as well as the average density of populations as compared to unstructured populations, see Fig. 4, and thus improves the odds of population survival.

Traditionally the effects of spatial structure in evolutionary games have been investigated based on lattices or more general network structures [23, 24]. Such discrete spatial arrangements are capable of supporting cooperation because they enable cooperators to form clusters and thereby reduce exploitation by defectors. This effect is further enhanced by success-driven migration [25]. In contrast, in our setup space is continuous and the state of the population represented by density distributions of cooperators and defectors. This difference is crucial and renders cooperation even more challenging because, in general, the density of defectors may become arbitrarily small but never and nowhere zero within finite times. As a consequence cooperators are always subject to exploitation by defectors. In fact, in the absence of ecological dynamics and constant population densities cooperators invariably disappear (if *r < N*) or, conversely, take over (if *r > N*), just as in well-mixed populations.

Notably, the only exception to this rule refers to the aggregation of cooperators because this mode of directed migration can trigger a positive feedback between densities and migration of cooperators: increases in density due to cooperator aggregation result in steeper gradients, which heightens the attraction and hence further increases their density. As a result, cooperator densities may not only remain localized but also eliminate defectors. However, this positive feedback results in analytical and numerical challenges because it gives rise to discontinuous distributions. Yet, those challenges are common among models incorporating aggregation. For example, the Keller-Segel chemotaxis model predicts infinite population densities after finite times [26–29]. Even though a mathematical artifact, those singularities are associated with the inherent feedback between chemotaxis and the secretion of chemical attractants [15].

Nevertheless, the most striking feature of directed migration is the potential of aggregating cooperators to crowd out and eliminate defectors altogether, see Fig. 2A, Fig. 3A,C, Fig. 4A. Unfortunately this intriguing phenomenon is linked to the positive feedback that gives rise to discontinuous distributions and hence eludes further analysis based on the present framework, Eq. (4). Somewhat surprisingly, the reduced aggregation rate due to a lack of reproductive opportunities, represented by the term *w∇u*, turns out to be insufficient to maintain smooth numerical solutions.

Motility plays an important role in biofilms. Microbes excrete extracellular substances to generate and maintain this protective film. Free-riders benefit from the protection without contributing, which gives rise to the public goods dilemma. Experiments indicate that not only the inherent social conflict plays a vital role in the effective secretion of biofilms [30] but also the microbes motility [31–33]. Biofilms are no longer generated when deactivating the movement apparatus through deliberate mutations [34]. When specifically targeting the ability to chemotact, biofilm production significantly varies across microbes and experimental setups [31]. This sensitivity of public goods production in response to different types of migration is reflected in our model.

The complex interplay between ecological public goods and motility shapes population densities and distributions as well as their social composition. Not only does migration affect the production of the public good but some public goods also alter the motility of their producers. For example biofilms increase viscosity and reduce the motility of microbes or even segregate populations [35]. Conversely, *Paenibacillus* collectively lubricate hard surfaces to enhance population expansion [36]. Either scenario creates intriguing opportunities for novel feedback mechanisms where migration not only shapes the production and availability of the public good but where the public good represents the very infrastructure needed to stay put or migrate more efficiently.

## Acknowledgments

We would like to thank Prof Michael Doebeli and Prof Joe Yuichiro Wakano for their helpful feedback and comments. Financial support is acknowledged from the Natural Sciences and Engineering Research Council of Canada (NSERC), grant RGPIN-2015-05795 (C.H.).

## Supplementary Information

**S1 Appendix. Derivation of directed migration terms.**

**S2 Appendix. Pattern formation: Dispersion relation and derivation of necessary criteria.**

**S1 Figure. Dispersion relation for directed migration.** (A) aggregating cooperators, *A*_*C*_, and (D) spreading defectors, *R*_*D*_, have both the potential to cause spatial instabilities through increased migration. In the process, the dominant mode *k_∗_* increases with increased aggregating cooperators, *A*_*C*_, and decreases with spreading defectors, *R*_*D*_. In contrast, (C) hunting defectors *A*_*D*_ and (D) fleeing cooperators *R*_*C*_ stabilize the system. *Parameters:* 2.4 = *r > r*_Hopf_ (such that *λ*(0) *<* 0), as well as *D_C_ > A_C_ u*_eq_*w*_eq_ (to ensure *λ*(*k*) *<* 0 for *k → ∞*). (A), (D) *D*_*D*_ = 0.5, (B),(C) *D*_*D*_ = 0.7. Other parameters as in Fig. 1.

**S2 Figure. Spatial and temporal dynamics for different types of directed migration.** Each panel illustrates the dynamics for a single type of directed migration as a function of the multiplication factor *r* in the proximity of *r*_Hopf_. The small rectangles depict the cross section of the density distribution through the middle of the square *L × L* domain as a function of time from top to bottom. The color brightness indicates the density of cooperation (green) and defection (red) with coexistence (yellow) and vacant space (black). (A) Aggregating cooperators oppose the effects of diffusion and hence slow down the population expansion. The positive feedback between cooperator densities and aggregation can cause discontinuous distributions and the breakdown of numerical methods. (B) Hunting defectors increase competition in areas where cooperators concentrate. This allows cooperators to escape into unpopulated terrain but also increases the risk of extinction. (C) Fleeing cooperators avoid defectors and hence readily explore vacant space but with the defectors at their tails also tend to spread themselves too thin and risk extinction. (D) Spreading defectors again promote pattern formation and thereby support the survival of the population. *Parameters:* same as in Fig. 1 but with Gaussian initial condition without noise and *t* = 250.

**S3 Figure. Pattern formation and discontinuities for aggregating cooperators** *A*_*C*_. Analytical findings distinguish three domains based on *A*_*C*_ and the multiplication factor *r*: (i) aggregation is sufficiently strong to destabilize the homogeneous equilibrium (grey shaded region, *A_C_ u*_eq_*w*_eq_ *> D_C_*); (ii) unstable modes give rise to pattern formation (red shaded region, see Eq. (S4) in S2 Appendix); (iii) stable homogeneous coexistence (unshaded region). However, initial conditions significantly impact the emerging dynamics. Heterogeneous initial distributions trigger migration and aggregation of cooperators. Discontinuities emerge within 250 generations from a Gaussian initial distribution when the selection-diffusion system is numerically integrated (above dashed horizontal line). Below the dashed horizontal-line, patterns emerge (in red shaded region) or homogeneous coexistence is regained (in unshaded region) as predicted. For small *r* heterogeneous distributions are unable to develop and the population goes extinct (left of dashed vertical line). *Parameters:* same as in S2 Fig, except *D*_*D*_ = 0.5 to promote smooth patterns.

